# *Pseudomonas aeruginosa* LasR-deficient mutants have increased methylglyoxal and hydrogen peroxide sensitivity due to low intracellular glutathione

**DOI:** 10.1101/2024.09.25.615034

**Authors:** Marina Ruzic, Ana I. Altamirano Hefferan, Amy Conaway, Deborah A. Hogan

**Affiliations:** Department of Microbiology and Immunology, Geisel School of Medicine at Dartmouth, Hanover, NH, USA

## Abstract

The electrophile methylglyoxal (MG) is produced by microorganisms and host cells through central metabolic pathways. MG is a highly reactive electrophile, so it must be rapidly detoxified to prevent damaging modifications to macromolecules. *Pseudomonas aeruginosa*, a pathogen of concern due to its ability develop multidrug resistance, causes many types of infections that have been associated with elevated MG levels, including cystic fibrosis (CF). *P. aeruginosa* isolates commonly have mutations that lead to LasR loss-of-function (LasR-) and we found that *lasR* mutations confer sensitivity to MG in multiple strain backgrounds. LasR-strains have increased activity of the CbrAB two-component system which represses Crc regulation of metabolism. Here, we show that higher CbrAB activity and low Crc activity renders cells sensitive to MG. We found that *P. aeruginosa* LasR-strains are more sensitive to MG and have lower intracellular reduced glutathione (GSH) compared to their LasR+ comparators. Consistent with published reports, mutants lacking *gloA3*, which encodes a MG-glyoxalase, and mutants lacking GSH biosynthesis enzymes (*gshA* or *gshB*) were sensitive to MG. Exogenous GSH rescued MG sensitivity in LasR-strains and *gshA* or *gshB* mutants, but not in a *gloA3* mutant strain. We propose that low GSH levels in LasR-strains contribute to increased sensitivity to MG and H_2_O_2_.

**Significance:** Methylglyoxal is a highly reactive metabolite that is detected in various disease states, including those where *Pseudomonas aeruginosa* is present and MG resistance requires the glutathione-dependent glyoxalase enzyme GloA3 enzyme. This study reveals that *P*.*aeruginosa* strains with LasR mutations, which are commonly found in clinical isolates, are more sensitive to methylglyoxal (MG) and hydrogen peroxide due to lower intracellular glutathione levels and high activity of the CbrAB-Crc regulatory pathway. This could be significant for understanding the selective pressures that drive *P. aeruginosa* evolution in infection sites, as well as a better understanding of LasR-strain metabolism in infections such as those associated with cystic fibrosis.

## Introduction

Methylglyoxal (MG) is a small, electrophilic, highly reactive dicarbonyl molecule that is endogenously formed in most living cells. MG spontaneously forms from a variety of metabolites such as glyceraldehyde-3-phosphate (G3P) and dihydroxyacetone phosphate (DHAP) which form during metabolic pathways such as those for the catabolism of sugars or glycerol (1). MG can be highly toxic since it can react with the amines on proteins, nucleic acids, and lipids to form damaging adducts. To prevent MG accumulation, cells contain an enzymatic detoxification system called the glyoxalase system, which metabolizes MG (1). The glyoxalase system consists of two major enzymes Glo1 and Glo2, and is dependent on reduced glutathione (GSH), an antioxidant molecule (1). Glo1 catalyzes the formation of S-D-lactoylglutathione from the spontaneously formed MG–GSH hemithioacetal and Glo2 converts S-D-lactoylglutathione into D-lactate as a final product (2). Other detoxification pathways in multiple organisms have also been described, such as GSH-independent glyoxalases (3) and NADPH-dependent MG reductases (4). Elevated MG occurs at sites of infections, cancer, diabetes, dementia, cystic fibrosis, kidney and liver disease (1, 5-11) due to increased glycolysis, which leads to an increase in the spontaneous formation of MG (1, 10), as well as a reduction in GSH levels or enzymes involved in MG catabolism (11-13). For example, cystic fibrosis (CF), a genetic disease caused by mutations in the CF transmembrane conductance regulator (CFTR) gene, is associated with low Glo1 and a GSH deficiency (11, 14).

*Pseudomonas aeruginosa* causes an array of infections, including respiratory infections, infections in chronic wounds, eye infections, ear infections, urinary tract infections, and blood infections, in part because of its ability to resist diverse stresses. *P. aeruginosa* has three genes encoding putative glyoxalase I enzymes, similar to those in eukaryotic cells. Two are nickel/cobalt activating enzymes, GloA1 and GloA2, and the last is a zinc-dependent enzyme, GloA3 (15). The detoxification of MG or other carbonyl-containing compounds by glyoxalase enzymes occurs via covalent modification to reduced glutathione (GSH), a thiol-containing tripeptide (16). GSH is synthesized by GshA, which forms γ-glutamyl-cysteine from glutamate and cysteine, and GshB which forms GSH through the addition of glycine (16). *P. aeruginosa, gshA* and *gshB* mutants have sensitivity to several different stressors, including antibiotics from multiple families, bleach, hydrogen peroxide (H_2_O_2_), and MG (17-21). Because *P. aeruginosa* has high inherent resistance to antibiotics, particularly in biofilms, and it can become multidrug resistant, alternative antimicrobials have been sought. One such alternative is manuka honey, a unique product derived from the nectar of the *Leptospermum scoparium* plant, gathered by bees. Manuka honey exhibits potent inhibitory effects against *P. aeruginosa* growth by depolarizing and permeabilizing the membrane, compromising the pathogen’s survival and proliferation (22). MG is one of the major antimicrobial components in manuka honey, and both manuka honey and MG lead to an upregulation of *gloA3* (22, 23).

*P. aeruginosa* populations evolve in reproducible ways across infected individuals over time. One common type of mutation that repeatedly arises is loss-of-function (LOF) mutations in the gene that encodes the transcription factor LasR, which participates in acylhomoserine-mediated quorum sensing (24). More recently, LasR has been shown to influence metabolism through the two-component system CbrAB and the downstream translational repressor Crc (25), which regulates target mRNAs involved in uptake of non-preferred carbon sources, stress response, and other metabolic processes (26-28). We previously provided strong evidence that LasR-strains have higher levels of CbrAB activity and lower levels of Crc-mediated repression of metabolism (25), and that these metabolic changes lead to the increased fitness of LasR-cells in passaged cultures (25). LasR-strains are commonly found in the environment and infection sites, and are of clinical relevance since they have been associated with worsened disease outcomes such as in CF (29). Interestingly, LasR-mutants have been found to be sensitive to oxidative stressors such as H_2_O_2_ as a result of less induction of catalases and NADPH-producing dehydrogenases (30). Though, the effects of MG on LasR-strains have not been previously reported.

In this study we found that *Pseudomonas aeruginosa* LasR-strains are more sensitive to MG than their LasR+ comparators, and genetic analyses indicate that high activity of the CbrAB two component system and low Crc repression of metabolism explains LasR-cell sensitivity to MG. We show that *gloA3* and GSH biosynthetic enzymes are essential for MG resistance and that GloA3-dependent MG resistance can be restored by exogenous GSH. GSH levels were significantly lower in LasR-strains, compared to their LasR+ counterparts. The previously published H_2_O_2_ susceptibility of LasR-strains was also found to be rescued by exogenous GSH. Together, these data suggest that LasR-strains have low intracellular GSH which can render them sensitive to MG and H_2_O_2_. These data may contribute to our understanding of why these LasR-strains frequently arise to predominance in CF lung infections but rarely persist under other growth conditions and over evolutionary time periods.

## Results

### *P. aeruginosa* strains lacking a functional *lasR* gene have increased sensitivity to extracellular MG

To determine the MG sensitivity of WT and Δ*lasR*, LB-grown cells were plated on LB containing MG at a range of concentrations (3-4mM). These concentrations were chosen based on our ability to detect differences in sensitivity between strains, as there is not a clinically relevant concentration due to the difficulty in measuring MG *in vivo. P. aeruginosa* strain PA14 wild type (WT) gave rise to ∼10-fold fewer colonies when plated on MG, and effects were dose-dependent (**Fig. 1A**). When compared to the WT, the Δ*lasR* mutant had significantly fewer CFUs on MG-containing media at all tested concentrations (**Fig. 1A**). Complementation of the Δ*lasR* strain (Δ*lasR + lasR*) restored MG resistance levels of the WT strain (**Fig. 1B**).

**Figure 1.**
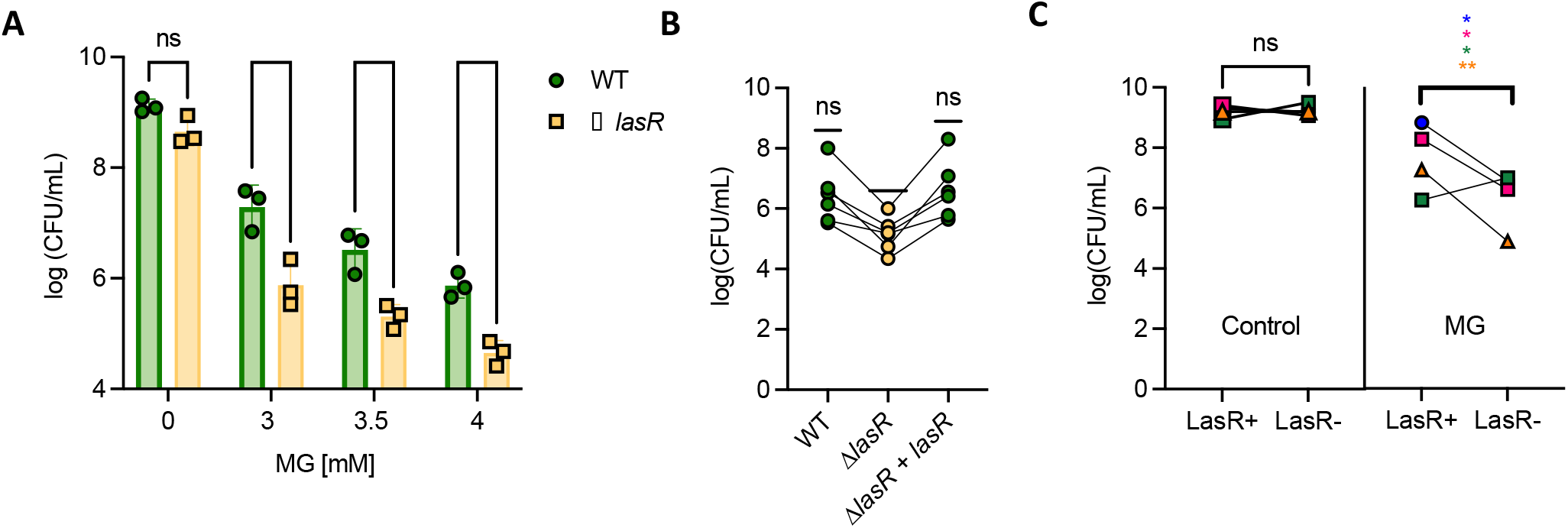
Mutations in the *lasR* gene result in sensitivity to methylglyoxal. (A) Methylglyoxal sensitivity assay of PA14 wild type (WT) and Δ*lasR* strains on LB agar with different concentrations of MG. Significance between WT and Δ*lasR* was determined with a paired t-test (n=3). (B) Methylglyoxal sensitivity assay at 3.5mM MG on WT, Δ*lasR*, and Δ*lasR + lasR*. A one sample t-test was performed, using WT mean as the theoretical mean (n=5). C) Methylglyoxal sensitivity assay at 3.5mM MG on four pairs of clinical isolates containing a functional and non-functional LasR. Significance was determined with a paired t-test. Different colors/shapes refer to different clinical isolate pairs (* indicates P<0.05, ** indicates P<0.01, *** indicates P<0.001, and **** indicates P<0.0001).

To further examine whether the absence of LasR activity increased MG sensitivity, we tested multiple clinical isolate pairs of LasR*+* (LasR functional) and LasR- (LasR non-functional) from the same clinical sample. The LasR+ and LasR-clinical isolate pairs displayed similar growth on LB agar. However, on LB agar with 3.5 mM MG, 3 out of 4 of the LasR-clinical isolates had less growth than LasR+ isolates. Together the LasR-strains were significantly more sensitive than their LasR+ counterparts (**Fig. 1C**).

### Genes involved in the CbrAB pathway play an important role in MG detoxification

LasR-strains have higher CbrAB pathway activity (25), thus, we sought to determine if differences in MG sensitivity between WT and Δ*lasR* were related to differences in CbrAB pathway activity (see **Fig. 2A** for pathway). We found that the Δ*crc* mutant had significantly worse growth on LB with 3.5 mM MG than the WT, and that its sensitivity to MG was similar to that of Δ*lasR*. The sensitivity of a double mutant Δ*lasRΔcrc* was higher than the single mutant (**Fig. 2B**).

**Figure 2.**
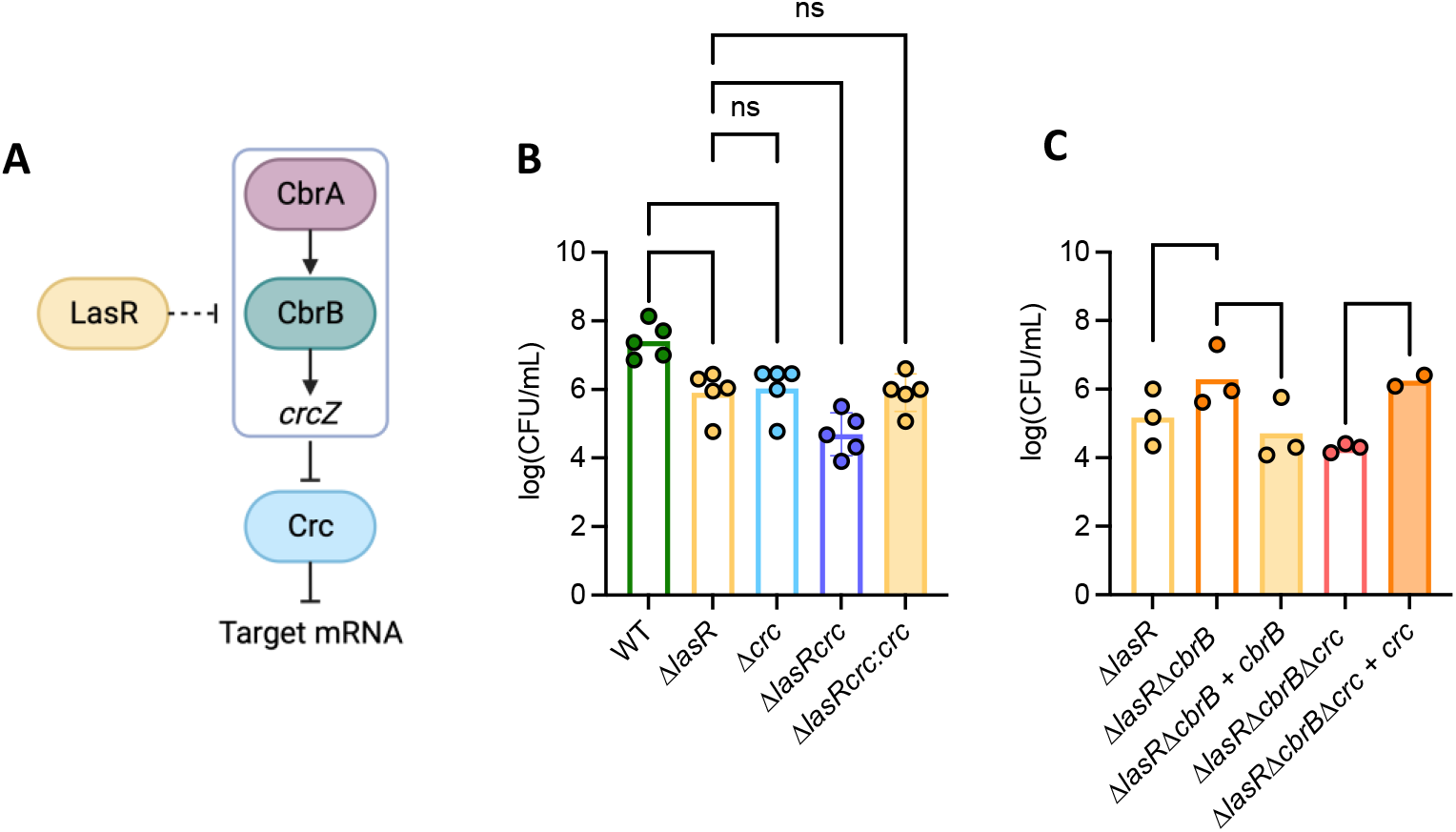
The deletion of genes in the CbrAB pathway influences sensitivity or resistance to methylglyoxal. (A) Carbon catabolite pathway: CbrAB induces expression of the small RNA *crcZ* which then sequesters Crc to allow translation of genes involved in metabolism. Previous evidence links LasR to the repression of CbrB, but its mechanism has not been characterized. (B-C) Methylglyoxal sensitivity assay at 3.5mM MG of knockout strains of genes within the CbrAB pathway. Complemented strains are shown next to their knock-out counterparts. Significance was calculated using a two-way ANOVA test (* indicates P<0.05, ** indicates P<0.01, *** indicates P<0.001, and **** indicates P<0.0001).

We found that the double mutant Δ*lasRΔcbrB* was significantly less sensitive than Δ*lasR* and that the complemented mutant Δ*lasRΔcbrB + cbrB* restored sensitivity of the Δ*lasR* strain to MG. A triple mutant Δ*lasRΔcbrBΔcrc* was more sensitive than Δ*lasR*, and the complemented Δ*lasRΔcbrBΔcrc* + *crc* strain had similar growth as Δ*lasRΔcbrB* (**Fig. 2C**).

### The overexpression of *gloA3* provides similar improvement in MG resistance across strains

*P. aeruginosa* had multiple candidate glyoxalases, but GloA3, a zinc-dependent glyoxalase I enzyme, has been specifically implicated in MG detoxification (15) (**Fig. 3A** for pathway). We tested growth of two transposon glyoxalase-encoding enzyme-producing genes, *gloA1*::*MAR2* and *gloA3*::*MAR2*, on 3.5mM MG (**Fig. 3B**). There was no significant difference between WT and *gloA1*::*MAR2* on LB with MG. Meanwhile, there was a large and significant difference between the growth of WT and *gloA3*::*MAR2*, indicating its importance in MG detoxification (**Fig. 3B**).

**Figure 3.**
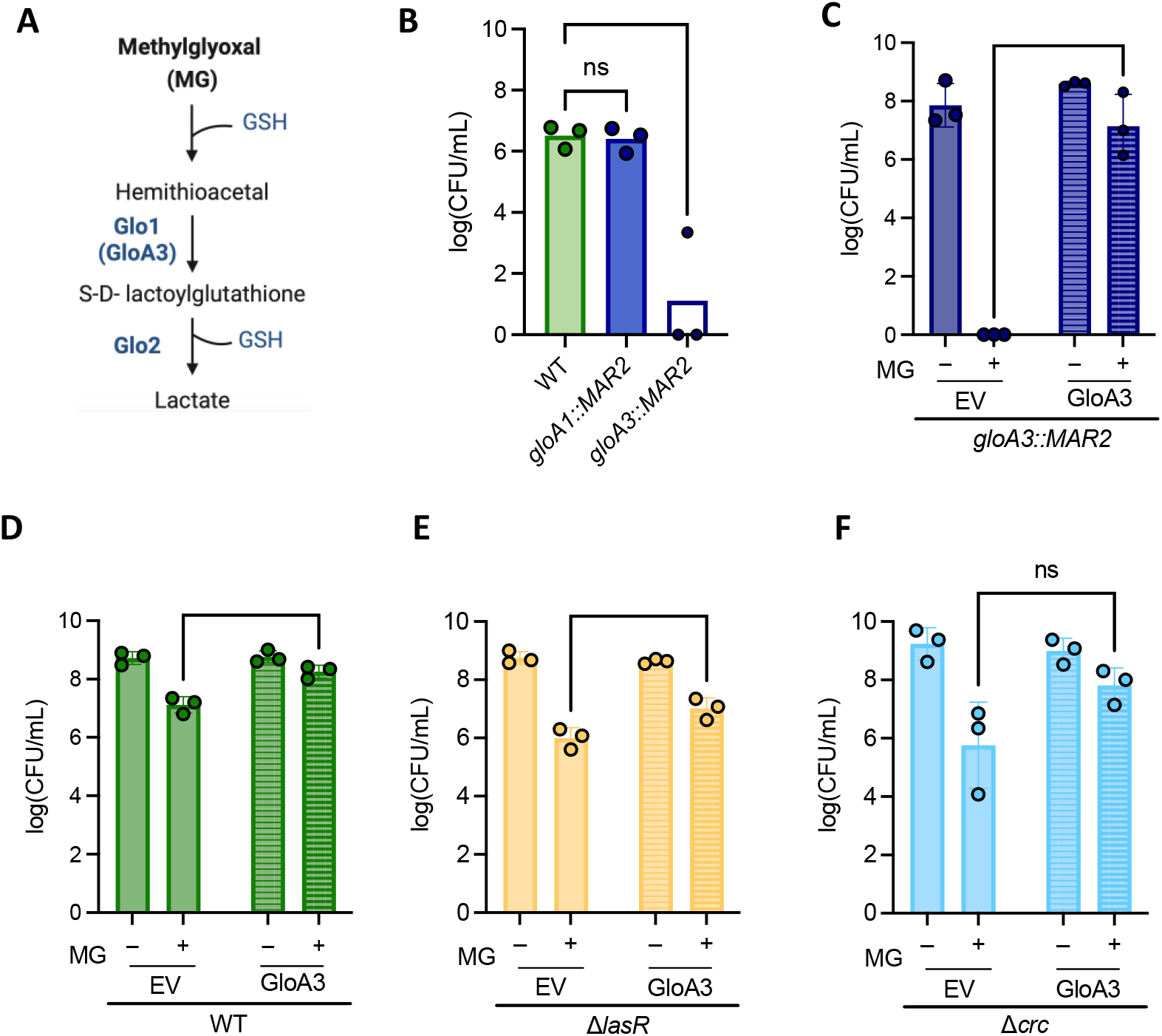
The gene *gloA3* is involved in protection against methylglyoxal. (A) Methylglyoxal detoxification pathway by glyoxalase enzymes (Glo1 and Glo2) and reduced glutathione (GSH) to yield D-Lactate. GloA3 is a Glo1 enzyme. (B) Methylglyoxal sensitivity assay at 3.5mM MG of transposon mutants *gloA1*::*MAR2* and *gloA3*::*MAR2* in relation to PA14 WT (n=3). Significance was calculated using a two-way ANOVA test. (P>0.01). (C-F) Methylglyoxal sensitivity assay at 3mM MG of strains containing a *gloA3* overexpression plasmid (GloA3) or an empty vector control (EV) (n=3). The following strains are shown in the graphs: (C) *gloA3* transposon mutant, (D) PA14 WT, (E) Δ*lasR*, and (F) Δ*crc*. A two-way ANOVA was performed to look at differences between strains or conditions (** indicates P < 0.01, *** indicates P < 0.001, and **** indicates P < 0.0001) The percent increase of p*gloA3*-containing strains compared to empty vector-containing strains is not significant between WT, Δ*lasR*, and Δ*crc*.

To perform complementation analyses, we introduced either a constitutively expressed *gloA3* on a plasmid or the empty vector pMQ70_EV (pEV), into the WT and *gloA3* mutant. Without MG, there was no significance in growth between *gloA3*::*MAR2* containing p*gloA3* or pEV (**Fig. 3C**), and the same trend was seen in WT (**Fig. 3D**). On medium with 3.5mM MG, the *gloA3*::*MAR2 +* pEV strain was not able to grow, while the p*gloA3* strain showed growth comparable to WT, showing that the constitutively expressed *gloA3* complemented the *gloA3::MAR2* mutant (**Fig. 3C**). The WT + p*gloA3* strain had a significant boost in growth when exposed to MG when compared to WT + pEV (**Fig. 3D**). The same trend was seen in Δ*lasR* + pEV and Δ*lasR* + p*gloA3* (**Fig. 3E**). The two Δ*crc* strains containing plasmids were not statistically significant from one another, however, Δ*crc* + *gloA3* appeared to have improved growth for each replicate in MG (**Fig. 3F**).

### Intracellular and extracellular GSH increases resistance to MG

It has previously been shown that *P. aeruginosa* strain PAO1 requires genes *gshA* and *gshB* for MG detoxification (21), and that GSH is necessary for GloA3 function (15). We aimed to determine if the same trend was seen in *P. aeruginosa* strain PA14, and whether intracellular GSH levels varied between WT and Δ*lasR* strains. In the presence of 3.5 mM MG, *gshA* and *gshB* transposon mutants (*gshA::MAR2* and *gshB::MAR2*) were unable to grow (**Fig. 4A**). On media containing 3.5mM MG and 5mM GSH, both *gshA::MAR2* and *gshB::MAR2* were greatly restored in growth. WT had improved growth when GSH was added to the media (**Fig. 4A**).

**Figure 4.**
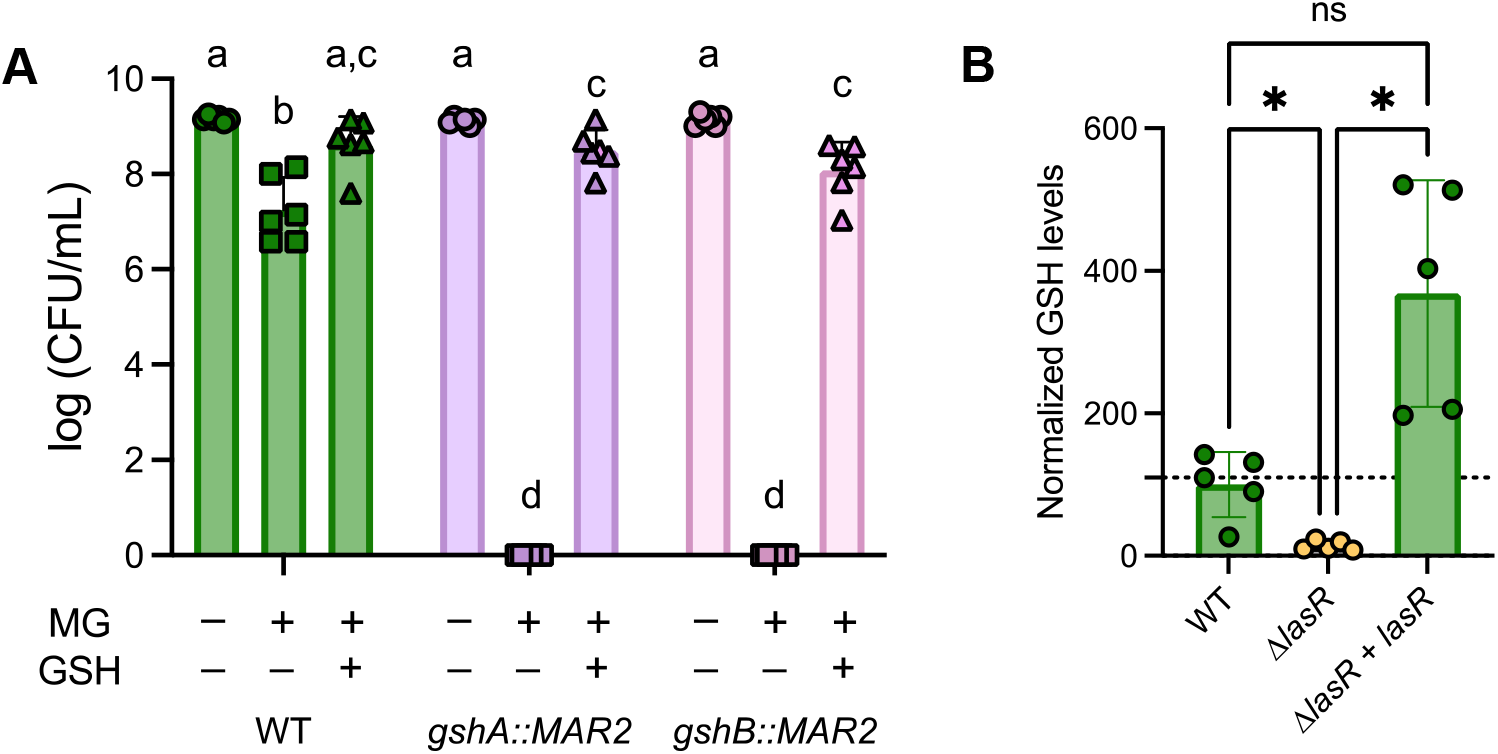
Glutathione is necessary for protection against methylglyoxal. (A) CFUs of WT and mutants lacking functional genes involved in glutathione biosynthesis (*gshA* and *gshB*) grown on LB agar alone or amended with 3.5 mM MG and 5 mM GSH as indicated. A two-way ANOVA was performed between and within media conditions. Bars with different letters are statistically significantly different (P<0.01, n=6). (B) Intracellular levels of glutathione were compared between WT, Δ*lasR*, and Δ*lasR* + *lasR* (n=5). Fold difference in amounts of glutathione were calculated, relative to wild-type. WT, Δ*lasR*, and Δ*lasR* + *lasR* significance is determined through a paired one-way ANOVA test (*, P < 0.05).

Endogenous levels of GSH were measured in WT, Δ*lasR*, and Δ*lasR + lasR* using a commercially available kit (GSH/GSSG ratio detection assay kit II, Abcam #ab205811). We found that Δ*lasR* had lower concentrations of GSH when compared to WT and its complement Δ*lasR + lasR* (**Fig. 4B**).

### Extracellular GSH protects against MG and H_2_O_2_ in different CbrAB backgrounds

Extracellular GSH was added to LB with and without 3.5mM MG (**Fig. 5A**) or 300μM H_2_O_2_ (**Fig. 5B**). Consistent with the results showing that Δ*lasR* sensitivity to MG is correlated with lower endogenous levels of GSH, the addition of GSH to LB + MG media partially rescued growth of Δ*lasR* significantly (**Fig. 5A**). In the WT, GSH completely rescued growth on medium with MG. For Δ*crc*, growth was improved when GSH was added to LB + MG media, but the difference between MG media with or without GSH was not significant. The transposon mutant strain *gloA3*::*MAR2* did not have significantly improved growth when GSH was added to media, suggesting that exogenous GSH provides protection against MG in a GloA3 dependent manner (**Fig. 5A**).

**Figure 5.**
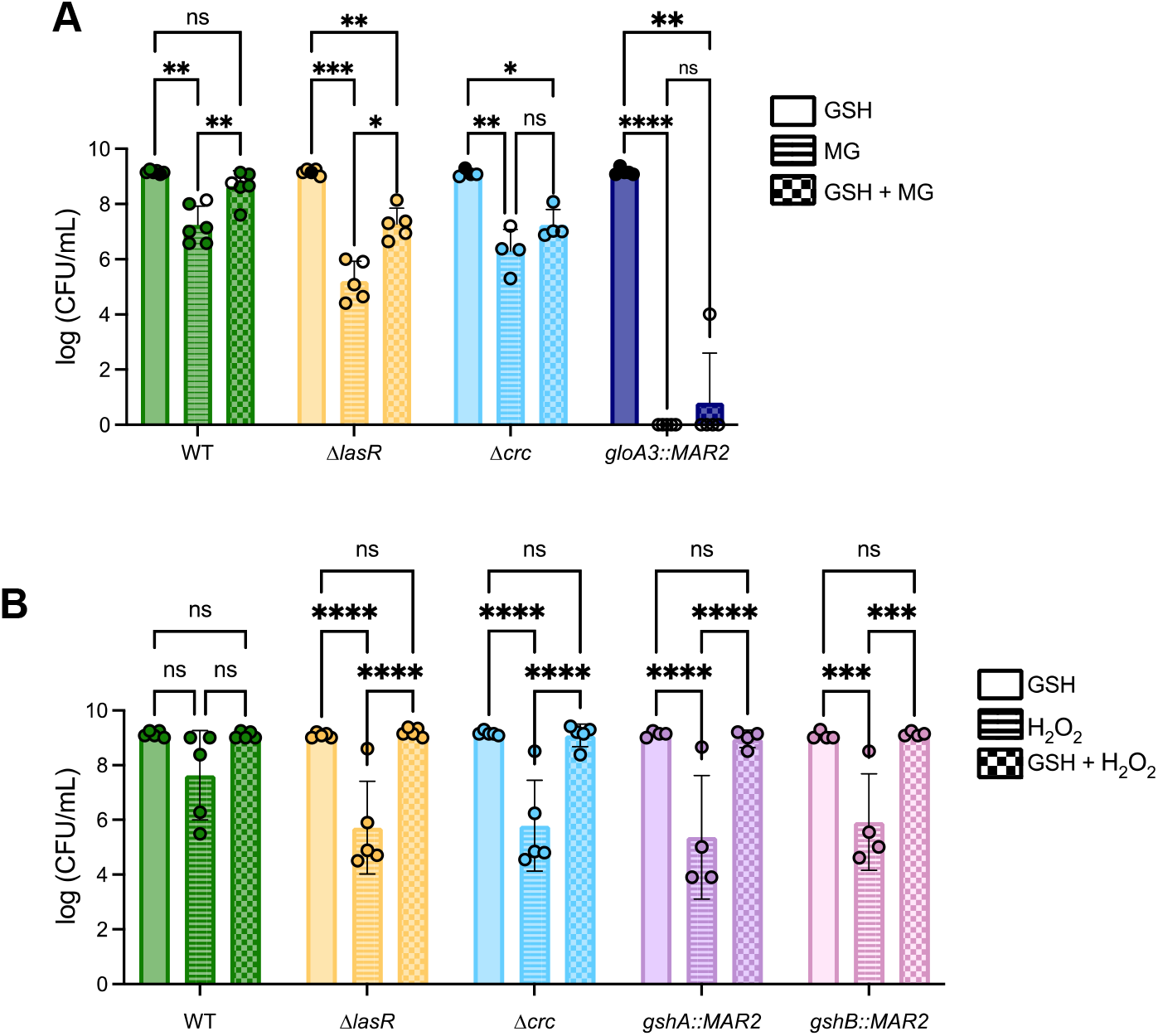
Extracellular glutathione protects against methylglyoxal and hydrogen peroxide in different strains. (A) Strain growth on LB agar containing 3.5mM MG compared to GSH and GSH/MG-containing agar (n=5). (B) Strain growth on LB agar media containing 300μM H_2_O_2_ is compared to media containing GSH and GSH/H_2_O_2_ (n=4). A two-way ANOVA was used to measure significance (* indicates P<0.05, ** indicates P<0.01, *** indicates P<0.001, and **** indicates P<0.0001).

It has been previously observed that the deletion of LasR and other quorum sensing genes increase sensitivity to H_2_O_2_ (31). H_2_O_2_ detoxification utilizes a pathway different from the glyoxalase pathway, using glutathione peroxidases (32-34), so we wanted to determine if GSH would also provide resistance to strains in the presence of H_2_O_2_ (**Fig. 5B**). When exposed to LB + 300 μM H_2_O_2_, *gshA::MAR2* and *gshB::MAR2* had significantly decreased growth when compared to media with no H_2_O_2_, but addition of GSH completely rescued growth. The Δ*lasR* and Δ*crc* mutants were significantly more sensitive than WT on H_2_O_2_ media, following the trend we had seen on MG. In all mutant strains, addition of GSH was able to completely rescue growth, pointing that GSH is an important molecule for protection against oxidative stress (**Fig. 5B**).

## Discussion

These studies lead us to propose a model in which the high sensitivity of LasR-strains to MG and H_2_O_2_ is due, at least in part, because of low intracellular GSH. Intracellular GSH concentrations were 8-fold lower in Δ*lasR* when compared to wild-type PA14 strain (**Fig. 4**). Overexpression of *gloA3* did not completely rescue differences between WT and Δ*lasR* strains grown in media with 3.5 mM MG (**Fig. 3**), but exogenous GSH did when GloA3 was present (**Fig. 5**). Our results also support our previous finding that the CbrA-CbrB-*crcZ*-Crc pathway is hyperactive in LasR-mutants (**Fig. 2A**) as deletion of *crc* increased MG sensitivity while deletion of *cbrB* increased resistance unless *crc* was already deleted (**Fig. 2 and Fig. S1**). Epistasis analyses also suggest that there are other effects of LasR loss-of-function that contribute to MG sensitivity (**Fig. 2B**).

The effects of MG on microbes in infections are not well understood. MG has antimicrobial properties, but microbes have mechanisms for MG detoxification. One study linked neutrophil-produced MG and *Streptococcus pyogenes* production of a MG-detoxifying glyoxalase I as important factors influencing outcomes in cell culture and animal model studies (35). MG also impairs control of *Mycobacterium tuberculosis* in infected bone-marrow derived macrophages (36). The differences in MG sensitivity between *P. aeruginosa* lab strains and clinical isolate pairs found in this study are similar to that of our previous studies with *Candida*, which described significant differences in MG susceptibility among *Candida lusitaniae* isolates that originated from a single patient infected only with this species (37). In that work, we found gain-of-function mutations in a gene encoding the transcription factor Mrr1 among *C. lusitaniae* CF clinical isolates (38) that led to the induction of MGD1, a MG reductase, and increased MG resistance as well as the increased capacity to use MG as a carbon source. Interestingly, MG induced Mrr1 regulation of other genes including *MDR1*, which leads to resistance to the antifungal fluconazole and other bacterial toxins (37). Thus, MG and changes to its transformation rate within microbial cells could influence the activity of MG-induced pathways. In fact, MG was shown to activate the CmrA pathway in *P. aeruginosa* leading to the upregulation of the efflux system MexEF-OprN, which can contribute to antibiotic resistance (39).

In *P. aeruginosa*, GSH, and thus the genes involved in GSH biosynthesis (*gshA* and *gshB*), are important for MG detoxification (17). Direct analysis of GSH levels in *P. aeruginosa* strain PA14 wild type and Δ*lasR* mutant found that *lasR* mutants had a significant reduction in GSH levels relative to PA14 WT when comparing average values (**Fig. 4B**). The less GSH available to act on MG, ultimately forming D-lactate as a less harmful byproduct, may explain the increased sensitivity. The lower levels of GSH in LasR-strains likely also explains the increased H_2_O_2_ susceptibility that has been previously documented (31). In addition to detoxification pathways, it has been shown that GSH also has a role in quorum sensing by integrating information about the redox state of the cell (40). Since LasR has a role in quorum sensing, it will be interesting to study the mechanisms which leads to lower GSH in LasR-mutants. GSH has multiple functions in the cell, as it can protect from different kinds of stresses such as nitrosative and oxidative stress, and even contribute to antibiotic resistance (41). GSH has also been shown to be involved in T3SS, swimming and swarming motility, and pyocyanin production (42, 43).

This study highlights the protective role of GSH against MG and H_2_O_2_ and shows the CbrAB two-component system as a contributor to MG sensitivity. We previously demonstrated that metabolic adaptations play a crucial role in the evolution and success of LasR-variants (25), and the results of this study point that MG and H_2_O_2_ sensitivity in addition to lower intracellular GSH of LasR-mutants could be contributing to this selection. Though, the sensitivity of LasR-mutants could change as we found one LasR-clinical sample isolate that was not sensitive to MG (**Fig. 1C**), which could be explained by *P. aeruginosa* evolution or upregulation of a GSH-independent mechanism. A GSH-independent glyoxalase (Glo3) has been characterized in *C. albicans, E. coli*, and *S. aureus* (44-46), but not in *P. aeruginosa*. Ultimately, the heterogeneity in MG detoxification mechanisms and CbrAB activity could contribute to the ability of *P. aeruginosa* strains to adapt in a wide array of environments.

## Materials and Methods

### Strains and growth conditions

*P. aeruginosa* strains used in these studies are listed in Table S1. Weekly, strains were struck out on Luria-Bertani (LB) agar plates (10 g/L peptone, 5 g/L yeast extract, 5 g/L NaCl (LB), 1.5% agar) from frozen stocks stored at -80°C. For overnight cultures, a single colony was transferred into 5 mL liquid LB medium in 18×150 mm glass tubes and incubated on a roller drum at 200 rpm for 16 h at 37ºC. The pMQ70-*gloA3* plasmid, synthesized by GenScript, or the pMQ70 empty vector control were electroporated into the indicated strain. Plasmid-containing cells were maintained on carbenicillin (300 μg/mL on agar plates and 150 μg/mL in liquid LB medium).

### Methylglyoxal and H_2_O_2_ sensitivity assays

Overnight cultures (∼16 h) grown in LB, adjusted to an OD_600_ of 1, then serially diluted by 10-fold dilutions in 200 μL of dH_2_O in a 96-well plate (Falcon). Five μL of each *P. aeruginosa* cell suspension dilution was spotted on the appropriate medium,dried in a biological safety cabinet for 30 min, then incubated for 16 h at 37ºC. The number of colonies from the last dilution with growth (greater than five colonies) were counted and each plate was imaged. LB agar amended with MG was prepared by adding the appropriate volume of MG (purchased as a 5.55 M liquid, Sigma M0252) to molten LB agar medium, then used within 3 h. The stated concentration of MG was added from a freshly prepared filter-sterilized 0.555 M stock solution in dH_2_O. H_2_O_2_ plates were prepared similarly, using a freshly prepared filter-sterilized 0.98 M stock (from a 30% w/w M H_2_O_2_ stock) added to the molten agar to a final concentration of 300 μM. For the GSH rescue experiments, reduced L-glutathione(Sigma #G2451) was diluted in water to a final concentration of 100 mM, filter sterilized, and added to molten LB agar to final concentration of 5 mM.

### Measurement of intracellular GSH

Cells were grown in 5 mL liquid LB at 37ºC on a rollerdrum for 16 h, then adjusted to an OD_600_ of 0.5. The GSH/GSSG ratio detection assay kit II (Abcam #ab205811) was used to measure intracellular GSH per manufacturer’s instructions. In brief, normalized cells were pelleted by centrifugation and washed with cold phosphate-buffered saline (PBS), then cells were lysed by treating with 0.5% NP40. Cells and debris were removed by centrifugation at 13,000 rpm for 15 min. The supernatant was treated with trichloroacetic acid (TCA) and NaHCO_3_ to remove proteins from the samples by incubating with each reagent for 5-10 min according to the manufacturer’s instructions. Eight GSH standards (0-10 μM) were prepared by diluting a 1mM GSH stock solution (1:100). To measure intracellular GSH, the GSH assay mixture was added to each sample and GSH standards, then incubated for 10-60 min. Fluorescence was measured at Ex/Em = 490/520 nm with a fluorescence microplate reader (BioTekSynergy Neo2). GSH of each sample was calculated by comparing fluorescence values to fluorescence obtained in the GSH standard curve.

### Statistical analysis and figure preparation

All graphs were prepared with GraphPad Prism 10.2.0 (GraphPad Software). One- and two-way analysis of variance (ANOVA) tests were performed in Prism; details on each test are described in the corresponding figure legends. All p values were two-tailed and p < 0.05 were considered to be significant for all analyses performed and are indicated with asterisks or letters in the text: *p<0.05, **p<0.01, ***p<0.001, ****p<0.0001. The CbrAB and glyoxalase pathway diagrams (Fig. 2A and 3A) were designed using BioRender (biorender.com).

## Acknowledgements

Research reported in this publication was supported by grants from the Cystic Fibrosis Foundation HOGAN19G0. Additional support came from NIGMS P20GM113132 through the Molecular Interactions and Imaging Core (MIIC), STANTO19R0 from the Cystic Fibrosis Foundation and NIDDK P30-DK117469 (Dartmouth Cystic Fibrosis Research Center). We would also like to thank Dr. Nicholas Jacobs for very helpful comments on the data and manuscript.

